# Single-cell RNA-seq reveals early heterogeneity during ageing in yeast

**DOI:** 10.1101/2020.09.04.282525

**Authors:** Yi Zhang, Jincheng Wang, Yuchen Sang, Shengxian Jin, Xuezheng Wang, Gajendra Kumar Azad, Mark A. McCormick, Brian K. Kennedy, Qing Li, Jianbin Wang, Xiannian Zhang, Yanyi Huang

## Abstract

The budding yeast *Saccharomyces cerevisiae* has relatively short lifespan and is genetically tractable, making it a widely used model organism in ageing research. Here, we carried out a systematic and quantitative investigation of yeast ageing with single-cell resolution through transcriptomic sequencing. We optimized a single-cell RNA sequencing (scRNA-seq) protocol to quantitatively study the whole transcriptome profiles of single yeast cells at different ages, finding increased cell-to-cell transcriptional variability during ageing. The single-cell transcriptome analysis also highlighted key biological processes or cellular components, including oxidation-reduction process, oxidative stress response (OSR), translation, ribosome biogenesis and mitochondrion that underlie ageing in yeast. Remarkably, we uncovered a molecular marker, *FIT3*, that was linked to mitochondrial DNA loss and indicated the early heterogeneity during ageing in yeast. We also analyzed the regulation of transcription factors and further characterized the distinctive temporal regulation of the OSR by *YAP1* and proteasome activity by *RPN4* during ageing in yeast. Overall, our data profoundly reveal early heterogeneity during ageing in yeast and shed light on the ageing dynamics at the single cell level.

## Introduction

It has been known for a long time that budding yeast *Saccharomyces cerevisiae* have limited division potential, only producing a finite number of daughter cells before death^1^. This phenomenon is defined as replicative ageing, and the number of daughter cells produced before death is defined as the replicative lifespan (RLS)^2^. Owing to its relatively short lifespan, detailed knowledge of its biology and its easy genetic manipulation, *S. cerevisiae* is regarded as an ideal model organism to study ageing^3^. Indeed, many ageing genes and signaling pathways initially found in yeast have also been later found to be conserved in other organisms, such as *C. elegans, M. musculus* and even humans^4^.

A dilemma of replicative ageing research in yeast exists between the rarity of old cells among an exponentially growing population either on a solid agar plate or in liquid media and the large number of pure old cells conventionally required for biochemical, genomic or transcriptomic analysis. To address this problem, several approaches have been developed to enrich old yeast cells, including magnetic sorting, elutriation, genetic programming and even computation^5-9^. However, these methods have yet to be successful at simultaneously ensuring both the quantity and purity of the isolated old yeast cells much less distinguishing old but living cells from dead ones. In addition, conventional bulk population analysis of ageing yeast cells may likely obscure some specific features within sub-populations due to the average effect^10^. Recent advances in microfluidics and single-cell imaging have revealed some phenotypic details of replicative ageing in yeast^11-14^; however, a systematic and quantitative investigation of yeast ageing at the single-cell and transcriptome level would be highly valuable.

Here, we developed a single-cell RNA-seq approach to study the replicative ageing of yeast and quantitatively assessed the heterogeneity between single yeast cells. Instead of partially purifying millions of old cells, exploiting single-cell technologies enabled us to obtain novel insights into yeast ageing from hundreds of single cells with precise ages. By profiling the transcriptomic landscapes of single yeast cells, we observed an increased cell-to-cell transcriptional variability and identified key functional biological processes or cellular components that were highly enriched during ageing. We also found early heterogeneity during ageing, indicated by a molecular marker of iron transport linked to mitochondrial DNA loss, and successfully characterized the distinctive temporal regulation of transcription between slow-dividing and fast-dividing age subgroups.

## Results

### Isolation of single yeast cells during ageing and scRNA-seq

Single yeast cells from isogenic populations ultimately have different lifespans. In fact, this is a universal phenomenon of ageing across species, albeit in different forms and ranges. And previous single-cell imaging data of replicative ageing in yeast have provided evidence of such heterogeneity. For example, when re-analyzing the single cell imaging data from the microfluidic-based yeast ageing studies^11,12^, we can observe that as early as 8 hr after birth, the distribution of generations of single yeast cells had already become dispersed, and the ranges of the distribution gradually increased at 12 hr and 16 hr after birth (Supplementary Fig. 1a), showing that some cells always divided more rapidly than others ever since early in life. These early-stage cell division dynamics in yeast seems closely associated with replicative age, with a positive correlation between the generations at early time points (8hr, 12hr, 16hr) after birth and the RLS (R=0.46, 0.64, 0.73; P=9.6×10^−5^, 7.7×10^−9^, 7.7×10^−9^; Supplementary Fig. 1b) at the single-cell level. This new finding is consistent with the previous report that the division time of single yeast cells early in life is negatively correlated with RLS, and the division time increases dramatically when approaching the end of life^11^. It was also reported previously that early in life, the gene expression level of *HSP104*, which encodes a molecular chaperone that maintains proteostasis in yeast, negatively correlates with RLS^11,12^. Accordingly, after re-analyzing the single cell imaging data^11,12^, we also observed a negative correlation between the generations at early time points during ageing and the *HSP104* gene expression level indicated by a GFP tag fused to this gene in single yeast cells (R=-0.43, −0.51, −0.56; P=2.8×10^−4^, 8.4×10^−6^, 7.8×10^−7^; Supplementary Fig. 1c). Collectively, these single-cell imaging data indicate an early heterogeneity of cell divisions during ageing in yeast, and that the division dynamics early in life can predict lifespan.

**Fig.1.**
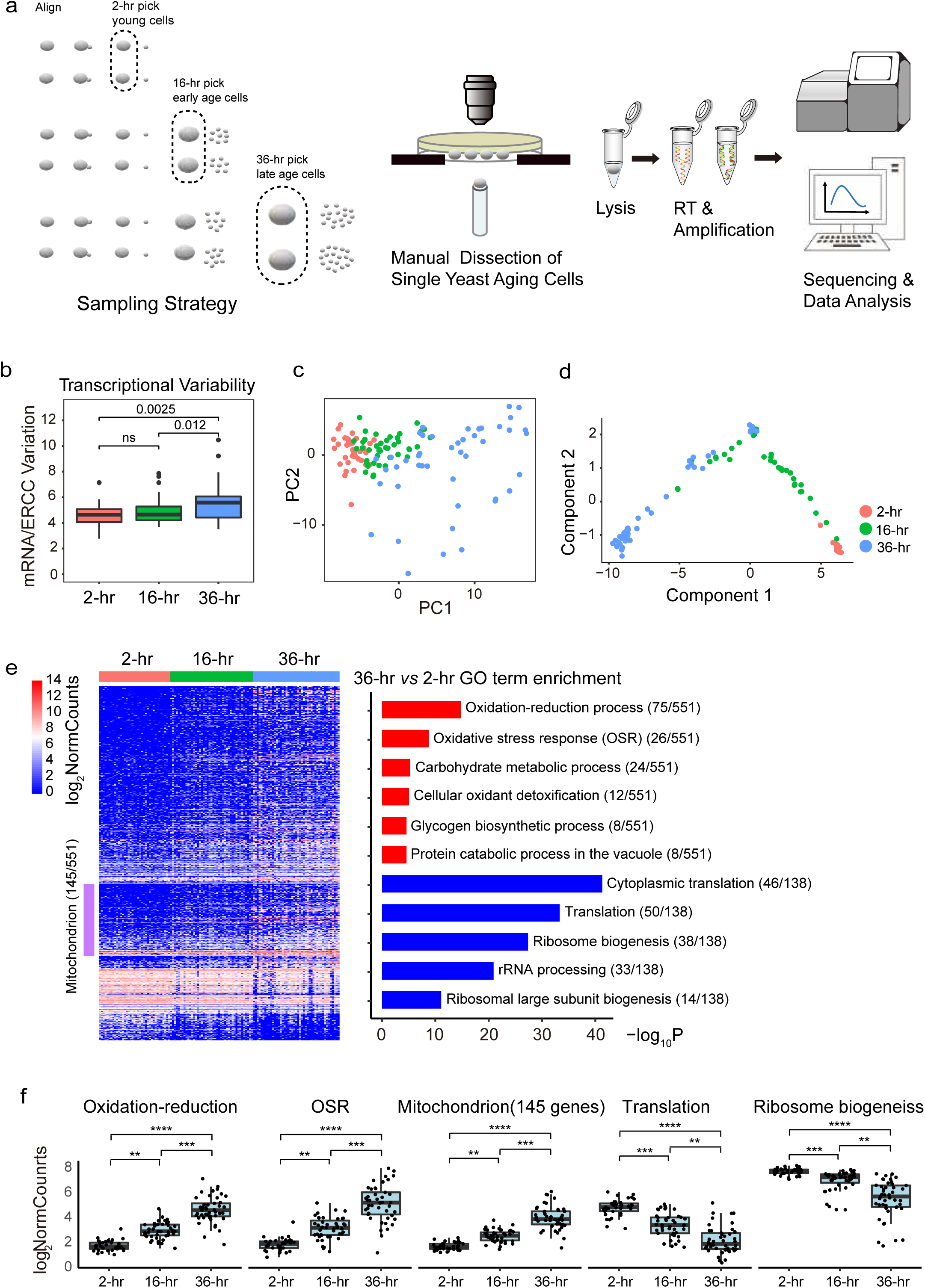
Cell-to-cell transcriptional variability and global differential gene expression during ageing in yeast. **a**, Schematic of the workflow of scRNA-seq during aging in yeast. Each single yeast ageing cell (indicated as gray ellipse in the dashed area) was manually isolated at 2-hr, 16-hr or 36-hr after birth, and then placed individually into a single tube prefilled with lysis buffer, followed by modified and optimized Smart-seq2^16,17^. **b**, Boxplot showing an increased cell-to-cell transcriptional variability during aging in yeast based on a correlation analysis where the transcriptional variability was measured as biological noise over the technical noise (see Methods). Boxes indicated the first and third quartiles, separated by median line. Whiskers indicated last values within 1.5 x the interquartile range for the box; Wilcoxon P values were also shown. **c**, PCA plot of single cells (n=125) from different age groups (no cell-cycle-regulated periodic genes included as input for PCA). The distribution of single yeast aging cells in the 36-hr late age group was more scattered than that of 2-hr age group and 16-hr early age group, which reflected an increased cell-to-cell transcriptional variability. **d**, Pseudotime trajectory of single cells (n=125) from different age groups (no cell-cycle-regulated periodic genes included as input). The youngest 2-hr age group was very concentrated, whereas the 16-hr early age group and 36-hr late age group were very scattered. **e**, (left) Heatmap of normalized gene expression of 551 upregulated and 138 downregulated genes in the 36-hr age group compared to 2-hr age group, across different age groups. The purple bar indicated 145 mitochondrial genes that were highly expressed in the 36-hr late age group. (right) Significance of GO terms of biological processes (BP) in upregulated and downregulated genes respectively (-log_10_P). **f**, Boxplot of the average normalized expression of significantly upregulated and downregulated gene categories identified in **e**, across different age groups. Each black dot in **f** represented a single cell. **p < 5.5 × 10^−7^, ***p < 4.2 × 10^−9^, ****p < 1.6 × 10^−13^, from Wilcoxon rank sum test.

To probe more deeply into the mechanisms underlying this early heterogeneity revealed by single-cell imaging, we further developed and applied scRNA-seq for transcriptome profiling of yeast during ageing (Fig. 1a; see Methods). We first conducted an RLS assay by continually performed manual microdissection of single yeasts on a solid agar plate^15^. At three different time points (2 hr, 16 hr and 36 hr after birth), we manually isolated single ageing yeast cells from the plate and placed the cells individually into a single tube prefilled with lysis buffer containing an external RNA control consortium (ERCC) spike-in for assessing technical noise then followed the Smart-seq2-based protocol^16,17^ with refined modifications and optimization for yeast ageing research (see Methods).

In total, we collected 136 yeast ageing cells for sequencing. The timepoints of isolation and number of generations at that time were precisely recorded for each cell (Supplementary Table. 1). After filtering out the cells with a low number of genes detected, insufficient read counts and ERCC-dominated samples (Supplementary Fig. 2a-c; see Methods), we finally retained scRNA-seq data of 125 cells composed of 37, 43 and 45 single cells in the 2-hr (young), 16-hr (early age) and 36-hr (late age) age groups, respectively, for further analysis. We also compared our scRNA-seq data to the only 2 available scRNA-seq datasets of *S. cerevisiae* published recently^18,19^. Our method yielded, on average, 2,202 genes detected per cell, which is comparable to the dataset from Gasch et al^18^ (2,202 *vs* 2,392) with good accuracy and sensitivity, similar to the dataset from Nadal-Ribelles et al^19^ (Supplementary Fig. 2d-e; Supplementary Table 1).

**Fig.2.**
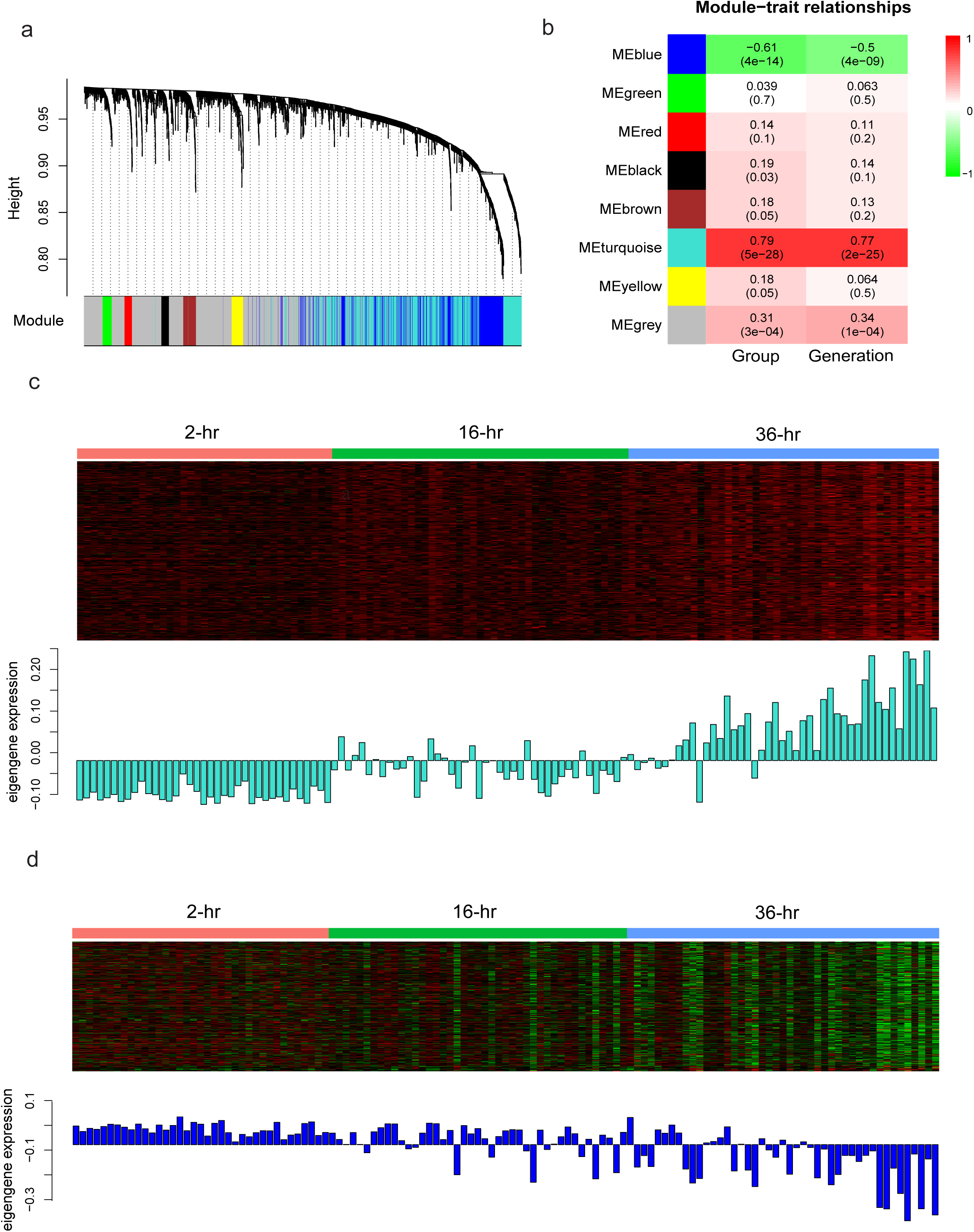
Weighted gene co-expression network analysis during ageing in yeast. **a**, Dendrogram showing the gene co-expression network constructed using WGCNA. The color bar labeled as “Module” beneath the dendrogram represents the module assignment of each gene. We totally identified 7 modules. **b**, Module-trait relationship shows that the turquoise module is most positively while the blue module is most negatively correlated with the traits of Group and Generation of the single yeast cells. The upper number within cell represents correlation coefficient and number within brackets refers to the p-value. **c** and **d**, Heatmap and barplot showing genes in the turquoise module are upregulated while genes in the blue module are downregulated during ageing in yeast. The rows of heatmap represent gene expression within the corresponding module. The columns of heatmap and barplot refer to the sample.

### Cell-to-cell transcriptional variability during ageing in yeast

We sought to explore the cell-to-cell transcriptional variability within different age groups using scRNA-seq data. Overall, we observed increased cell-to-cell transcriptional variability during ageing in yeast based on a correlation analysis in which the transcriptional variability was measured as the biological noise over the technical noise^20^ (Fig. 1b; see Methods). We verified this increase in cell-to-cell transcriptional variability alternatively using a quantitative statistical method^21^ and respectively identified 145, 312 and 524 highly variable genes (HVGs) with coefficients of variation (CV) that were significantly higher than those of the ERCC spike-in reference within each age group (Supplementary Fig. 3a; see Methods). Interestingly, by Gene Ontology (GO) analysis of these HVGs using DAVID^22^, the biological processes of cellular iron ion homeostasis and siderophore transport were specifically found to be highly enriched in the 16-hr early age group with high statistical significance, implying an early heterogeneity during ageing in yeast with regard to iron transport (Supplementary Table. 2).

**Fig.3.**
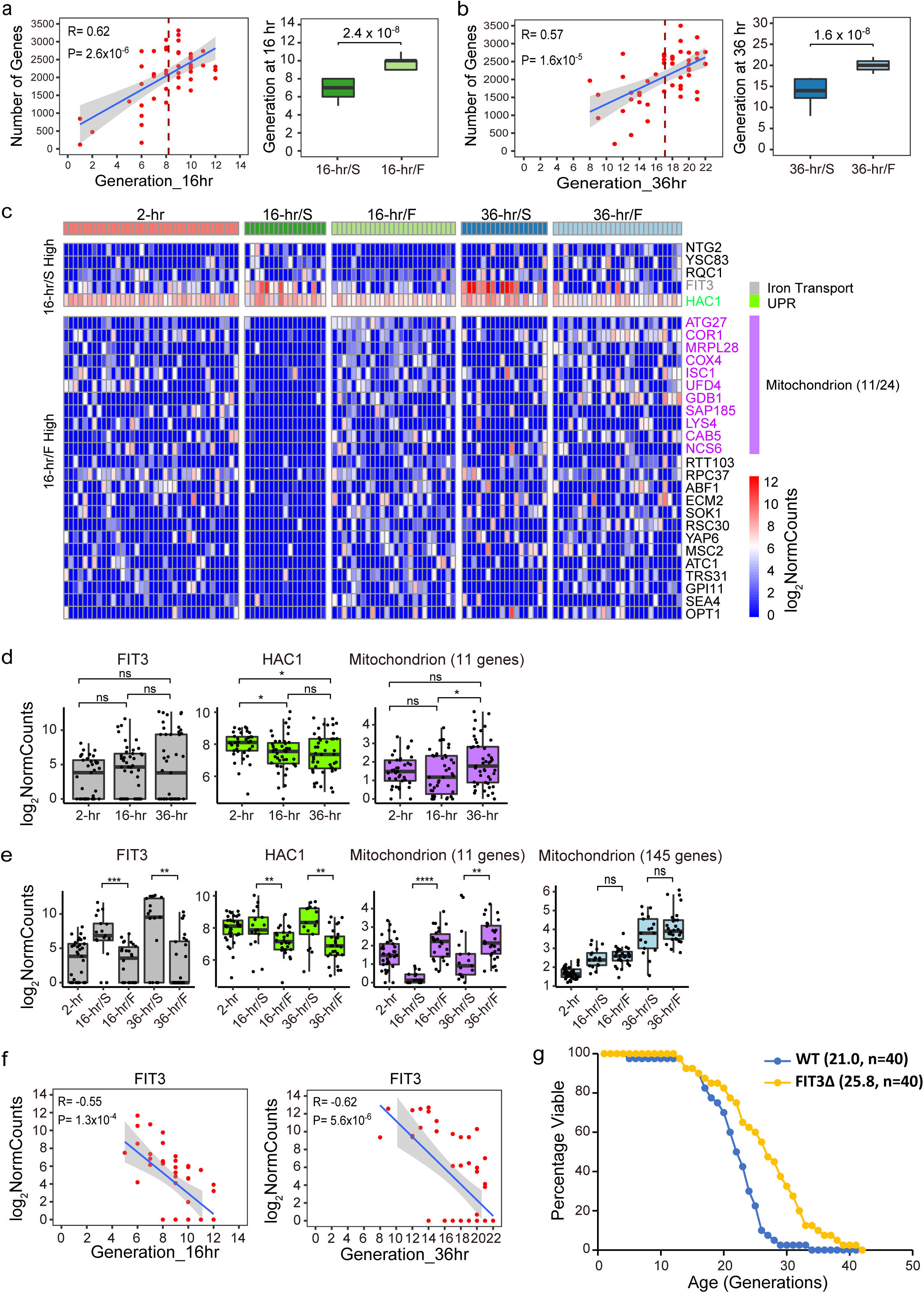
Differential gene expression between slow- and fast-dividing age subgroups. **a**, (left) Correlation of the number of genes detected and the generation of single cells in the 16-hr early age group. Each red dot represented a single cell with the number of genes detected and its generation at 16 hr. Blue line was a linear fit with gray area indicating 0.95 confidence interval; correlation coefficient (R) and P value (P) were also shown. The dashed line indicated the mean generation. The plot showed a positive correlation between the number of genes detected and the generation at 16 hr among individual cells. (right) Boxplot of generation between early age subgroups 16-hr/S and 16-hr/F that were split by the mean generation of 16-hr early age group; Wilcoxon P value was shown. **b**, (left) Correlation of the number of genes detected and the generation of single cells at 36-hr late age group and (right) Boxplot of generation between late age subgroups 36-hr/S and 36-hr/F that were split by the mean generation of 36-hr late age group, plotted same as in **a**. Note: The cells with the number of genes below 1000 plotted in both **a** and **b** were discarded in the rest analysis. **c**, Differential gene expression analysis between early age subgroups 16-hr/S and 16-hr/F. The heatmap showed normalized gene expression of statistically significant (Log_2_|FC|>1 and P_adj_<0.05) upregulated and downregulated genes in early age subgroup 16-hr/S compared to 16-hr/F, across different age subgroups. **d**, Boxplots of normalized expression of significant differentially expressed genes of *FIT3, HAC1*, and gene category of mitochondrion identified in **c** across different age groups. **e**, Boxplots of normalized expression of significant differentially expressed genes of *FIT3, HAC1*, and gene categories of mitochondrion respectively identified in **c** and **Fig. 1e** across different age subgroups. Each black dot in **d** and **e** represented a single cell. *p and **p < 0.05, ***p < 0.01, ****p < 6.1 × 10^−5^, and “ns” means not significant, from Wilcoxon rank sum test. **f**, Correlation of normalized gene expression of *FIT3* and the generation of single cells in the 16-hr early age group and 36-hr late age group, respectively. Each red dot represented a single cell. Blue line was a linear fit with gray area indicating 0.95 confidence interval; correlation coefficient (R) and P value (P) were also shown. The plot showed a negative correlation for both age groups. **g**, Survival curve of WT and *FIT3****Δ***. The number in the parenthesis represented the mean RLS and “n” indicated the number of cells assayed for RLS of each strain.

Because all of the ageing single yeast cells analyzed did not have synchronized cell cycles, we wondered whether and to what extent the cell-to-cell transcriptional variability was associated with the cell cycle. We found that 19.3%, 12.8% and 15.5% of HVGs, respectively, among the 3 age groups were regarded as cell-cycle-regulated periodic genes^23^ (Supplementary Fig. 3b). These results are consistent with a recent report of scRNA-seq in budding yeast that cell-cycle-regulated periodic genes were enriched in HVGs^19^. However, the trend of increased cell-to-cell transcriptional variability during ageing remained even when these cell-cycle-regulated periodic HVGs were removed from the 3 age groups (117, 272 and 443 HVGs remained, respectively; Supplementary Fig. 3b). We further confirmed this trend using principal component analysis (PCA). Regardless of whether the cell-cycle-regulated periodic genes were included in the scRNA-seq dataset used as input for the PCA or not, the 3 age groups were always successfully separated along the axis of first PCA component and were increasingly dispersed (Fig. 1c; Supplementary Fig. 3c); moreover, the top 30 genes based on the absolute loading values for the first PCA component always highly overlapped and were enriched in the biological process of cellular response to oxidative stress, which reflects ageing itself rather than the cell cycle (Supplementary Fig. 3d-e; Supplementary Table. 3). We also performed pseudotime analysis using Monocle^24^ and found that while the young cells (2-hr) were still very concentrated, the cells of the early age group (16-hr) had already become scattered along the trajectory (Fig. 1d; Supplementary Fig. 3f).

### Global differential gene expression during ageing in yeast

In addition to exploring the cell-to-cell transcriptional variability within different age groups, the scRNA-seq data also allow us to globally investigate the differential gene expression between age groups. Thus, we conducted a pairwise comparison among the 3 age groups using DESeq2^25^ (Supplementary Fig. 4a; see Methods). Obviously, more differentially expressed genes were found in the 36-hr late age group compared to the 2-hr group (Supplementary Fig. 4a, right panel; Supplementary Table. 4). The biological processes of oxidation-reduction and the oxidative stress response (OSR) were highly enriched in the 36-hr group (75 and 26 out of 551 genes, respectively), while translation and ribosome biogenesis were highly enriched in the 2-hr group (50 and 38 out of 138 genes, respectively) based on the GO analysis of biological process using DAVID^22^ (Fig. 1e, right panel). Moreover, 145 out of 551 genes that were highly expressed in the 36-hr late age group compared to the 2-hr group were enriched in mitochondrion as revealed by the GO analysis of cellular components (Fig. 1e, left panel; Supplementary Table. 4).

**Fig.4.**
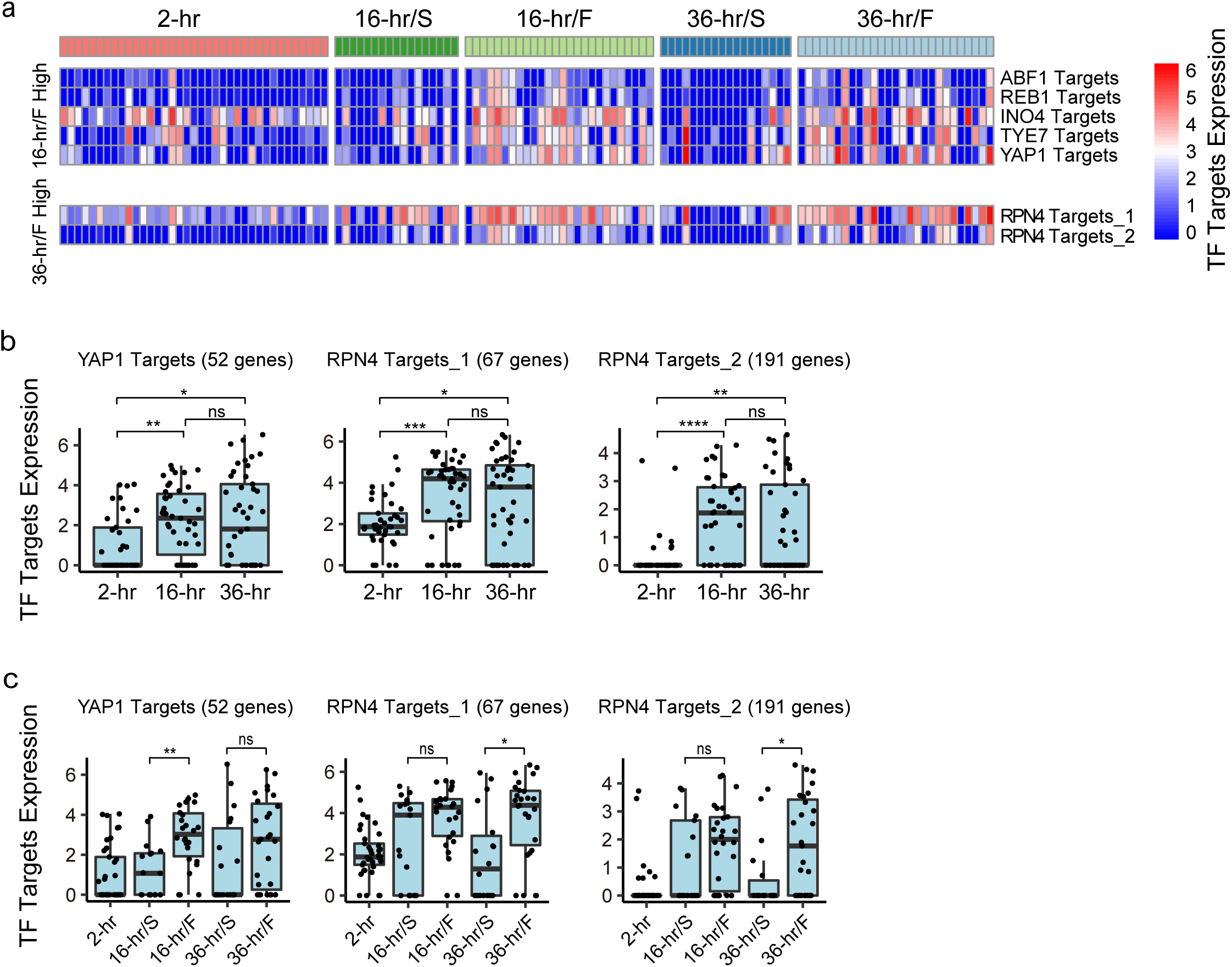
Temporal regulation of transcription factor (TF) between age subgroups. **a**, Heatmap showing differential expression of 5 transcription factor targets in the early age subgroup of 16-hr/F compared to 16-hr/S, and 2 transcription factor targets in the late age subgroup of 36-hr/F compared to 36-hr/S, based on first two statistical criteria (see Methods). **b** and **c**, Boxplots of differential expression of *YAP1* targets that were highly expressed in the early age subgroup of 16-hr/F compared to 16-hr/S, and 2 *RPN4* targets that were highly expressed in the late age subgroup of 36-hr/F compared to 36-hr/S identified by 3 stringent statistical approaches (see Methods), across different age groups and subgroups, respectively. Each black dot in **b** and **c** represented a single cell. *p < 0.05, **p < 0.01, ***p < 1 × 10^−3^, ****p < 1 × 10^−4^, and “ns” means not significant, from Wilcoxon rank sum test.

The average normalized gene expression levels across age groups further demonstrated an age-dependent increase in oxidation-reduction, OSR and mitochondrion as well as a decrease in translation and ribosome biogenesis (Fig. 1f). Indeed, these transcriptome changes had already occurred in the 16-hr early age group. Although far fewer differentially expressed genes were found in the 16-hr early age group compared to the 2-hr group (Supplementary Fig. 4a, left panel), early signs of upregulation in oxidation-reduction and downregulation in ribosome biogenesis (15 out of 108 genes and 4 out of 10 genes, respectively) were observed (Supplementary Fig. 4b; Supplementary Table 4). Notably, the global differentially expressed genes between age groups and their enriched GO categories from our scRNA-seq data were found to coincide well with a recent report of transcriptome changes during ageing in yeast^9^ and were even partially consistent with another proteome analysis of ageing in *C. elegans*^26^, although they were both based on bulk population analysis. These ageing associated GO categories analyzed by DAVID were also confirmed by ClusterProfiler^27^ (Supplementary Fig. 5a-f).

### Weighted gene co-expression network analysis during ageing in yeast

To find the clusters of highly correlated genes during ageing in yeast, we performed a weighted gene co-expression network analysis (WGCNA)^28-29^, and generated 7 different gene co-expression modules (Fig. 2a; see Methods). Among these gene co-expression modules (Fig. 2b-d), we further identified 52 hub genes from 731 genes in the positively correlated modules which were upregulated during ageing (Supplementary Table. 5). These genes are mainly enriched in OSR and oxidation-reduction process by GO analysis using Metascape^30^, and 5 of them are even involved in the longevity regulatory pathways, including *HSP104*, which is a molecular marker of ageing in yeast identified previously^11,12^ (Supplementary Fig. 6a; Supplementary Table. 5). 70 hub genes were identified from 410 genes in the negatively correlated modules which were downregulated during ageing and they are mainly enriched in ribosome biogenesis (Supplementary Fig. 6b). All these findings echo well the results of previous global differential gene expression analysis during ageing in yeast.

### Differential gene expression between slow- and fast-dividing age subgroups

The number of genes detected per cell within age groups was found to be positively correlated with the generation, suggesting another facet to understand the heterogeneity of cell divisions during ageing in yeast, and the 16-hr and 36-hr age groups were thus split by their respective mean generation into slow-dividing (16-hr/S, 36-hr/S) and fast-dividing (16-hr/F, 36-hr/F) subgroups (Fig. 3a, b; Supplementary Table. 1). Comparing the early age subgroups of 16-hr/S and 16-hr/F by DESeq2^25^ with stringent statistical filtering yielded 29 differentially expressed genes, with 5 highly expressed and 24 weakly expressed in 16-hr/S (Fig. 3c; Supplementary Table. 6). *FIT3* and *HAC1* are highly expressed in 16-hr/S. *FIT3*, together with *FIT2* and *FIT1*, as facilitators of iron transport in yeast, encodes a cell wall mannoprotein^31^. These genes were reported to be induced upon iron deprivation or mitochondrial DNA loss^32,33^. *HAC1* is a transcription factor that regulates the unfolded protein response (UPR), and interestingly, one of its regulatory targets is *FIT3*^34,35^. Indeed, *FIT3* and *HAC1* were not only highly expressed in 16-hr/S but also in 36-hr/S (Fig. 3d, e). Moreover, the gene expression of *FIT3* and *HAC1* negatively correlated with the age of single cells in the 16-hr age group (R=-0.55, -0.38; P=1.3 × 10^−4^, 1.5 ×10^−2^) as well as the 36-hr group (R=-0.62, -0.44; P=5.6 × 10^−6^, 2.2 × 10^−3^; Fig. 3f; Supplementary Fig. 7a; Supplementary Table. 6). Surprisingly, gene expression levels of several other iron transporters, including *FIT2* and *FET3*^31^, were also found to be negatively correlated with the generation of single cells in the 16-hr and 36-hr age groups (Supplementary Fig. 7b, c; Supplementary Table. 6). Finally, as single-gene deletions of *FIT2* and *FET3* were both reported to extend the lifespan in yeast^4^, we measured the RLS of yeast after deleting FIT3, and verified that this strain is long-lived as well (Fig. 3g). Collectively, these results clearly reveal a molecular marker of iron transport that can quantitatively indicate early heterogeneity during ageing in yeast, which might be mediated by mitochondrial DNA loss^33^. This early ageing transcriptional signature can last until an advanced age and predict the lifespan.

Interestingly, we also revealed that 11 out of 24 genes expressed at low levels in 16-hr/S were enriched in mitochondrion, and these genes were also expressed at lower levels in 36-hr/S than in 36-hr/F (Fig. 3c-e; Supplementary Table. 6). This further suggests a relatively poor mitochondrial function in the slow-dividing cells. Among these 11 weakly expressed mitochondrial genes (Fig. 3c), *COR1* is the core subunit of ubiquinol-cytochrome c reductase which belongs to complexes III and *COX4* is an important component of cytochrome c oxidase which belongs to complexes IV of the mitochondrial inner membrane electron transport chain. It has been reported that mutation of either *COR1* or *COX4* can cause a decrease in respiration, slow cell growth and even a shorter lifespan^34-38^. These 11 mitochondrial genes showed no overlap with the 145 mitochondrial genes that were globally upregulated during ageing (Fig. 1e and Fig. 3c, Supplementary Table. 4 and 6); in contrast, no significant differential expression of those 145 mitochondrial genes was observed between these two subgroups (Fig. 3e). These results successfully characterize divergent mitochondrial gene expression profiles between age groups and subgroups that would be masked in the bulk population analysis but can be identified by scRNA-seq.

The correlation analysis between the gene expression and the generation of single cells also resulted in genes that were positively correlated with generation in the 16-hr early age group are enriched in ribosome biogenesis (Supplementary Fig. 7d; Supplementary Table. 6). This suggests a downregulation of at least some ribosome biogenesis genes during early ageing and it is mainly contributed by the cells from the slow-dividing age subgroup, which are inclined to be short-lived (Supplementary Fig. 7e). Meanwhile, genes enriched in translation, mitochondrial translation and glycolytic processes were positively correlated with generation in the 36-hr late age group (Supplementary Fig. 7f). This agrees with the differential gene expression analysis above, suggesting a relatively poor machinery of translation and mitochondrion in the slow-dividing age subgroups. In summary, all these results thoroughly characterize early and late heterogeneity during ageing in yeast at the single-cell transcriptome level.

### Temporal regulation of transcription factor (TF) between age subgroups

We further investigated the regulatory variation in transcription factors (TFs) between age subgroups, analyzing 634 overlapping TF targets (gene clusters) based on reported studies on budding yeast^18,39-43^. To eliminate false positives, we performed stringent statistical analysis with three approaches (see Methods). First, we conventionally compared the median TF target expressions between age subgroups. This led to 16 TF targets that were significantly activated in the 16-hr/F subgroups and 11 TF targets in 36-hr/F compared to their counterparts, respectively (Supplementary Fig. 8a, b; Supplementary Table. 7). Then, we ran a Wilcoxon rank sum test comparing normalized gene expression levels of each set of TF targets to that of all other detected genes for each cell, taking P < 0.0001 as the criterion, followed by intersection with TF targets derived from the conventional analysis. This led to 5 and 2 TF targets that were significantly activated in 16-hr/F and 36-hr/F, respectively (Fig. 4a; Supplementary Fig. 8c; Supplementary Table. 7). Subsequently, we employed correlation analysis between TF target expression and the generation of single cells in the 16-hr and 36-hr age groups, taking P < 0.05 as the criterion (Supplementary Fig. 9a, b; Supplementary Table. 7), followed by intersection with TF targets derived from the former two approaches.

Finally, *YAP1* was found to be most significantly active in regulating the early age subgroup of 16-hr/F compared to 16-hr/S (Fig. 4b, c), although the other 4 TFs of *ABF1, REB1, INO4* and *TYE7* demonstrated a similar trend with less statistical significance (Supplementary Fig. 8d, e). Moreover, 2 TF targets of *RPN4* were found to be most highly regulated at 36-hr/F compared to 36-hr/S (Fig. 4b, c). *YAP1* is involved in activating the transcription of antioxidant genes in response to oxidative stress^44,45^. The relatively high activation of *YAP1* targets in the 16-hr/F early age subgroup suggests that the rapidly dividing single cells, which are inclined to be long-lived, may have a better defence system against oxidative stress than the slow-dividing cells. *RPN4* is a TF that stimulates proteasome biogenesis for the degradation of damaged proteins^46^. The relatively high activation of *RPN4* targets in the 36-hr/F late age but rapidly dividing subgroup supports the idea that proteasome capacity is critical to maintain the vigour and proteostasis of yeast cells, especially when approaching the end of life, as elevated *RPN4* expression is essential for extending the RLS in yeast^47^. Altogether, these findings reveal early and late heterogeneity by distinctive temporal regulation of TFs during ageing in yeast, and combined with the aforementioned differential gene expression analysis between age groups and subgroups, we successfully depicted a landscape of ageing in yeast with unprecedented detail at single-cell resolution.

## Discussion

Although transcriptome changes during ageing in yeast based on bulk population analyses have been reported^8,9^, such analyses at the single-cell level had not yet been performed. Here, we first identified an early heterogeneity of cell divisions during ageing in yeast by single-cell imaging and then developed and applied scRNA-seq for single-cell transcriptome analysis during ageing in yeast for the first time.

Using scRNA-seq technology, we overcame the difficulty of purifying the large number of old cells required for conventional transcriptome analysis during ageing in yeast. More importantly, by single-cell transcriptome analysis, we not only successfully recapitulated the results of the bulk population analysis but also teased out specific transcriptional features at the single-cell resolution that would otherwise be masked in a bulk population. For example, by scRNA-seq we revealed that while globally there were an age-dependent upregulation of many mitochondrial genes between age groups, a small number of different but important mitochondrial genes were significantly downregulated in the slow-dividing age subgroups compared to their fast-dividing counterparts. This provides novel and unprecedented insights into our understanding of the ageing process. Our results have unveiled the increased cell-to-cell transcriptional variability independent of the cell cycle and identified an early heterogeneity during ageing in yeast. This also coincides with recent reports of scRNA-seq in mouse immune cells and human pancreatic cells during ageing^48,20^.

By single-cell transcriptome analysis, we also identified a new molecular marker of iron transport that both indicates early heterogeneity during ageing in yeast and predicts lifespan. Remarkably, *FIT3* together with several other iron transporter genes, such as *FIT2* and *FET3*, had a negative correlation with the age of single yeast cells from both early and late timepoints. These genes are known to be induced upon iron deprivation or mitochondrial DNA loss^32,33^. Moreover, these genes can all extend the RLS in yeast when deleted^4^ (Fig. 3g). Therefore, we propose a model in which early heterogeneity during ageing in yeast is associated with differential mitochondrial dysfunction that affects and is mediated by iron transport. And this model is partially supported by a report published recently, showing age-dependent heterogeneity via a *FIT2* reporter that is correlated with vacuolar pH, mitochondrial function and lifespan in sub-populations of yeast cells^50^. More evidence may be needed to further validate this model, and presently it remains challenging to disentangle the cause-effect relationships between mitochondrial dysfunction and early heterogeneity during ageing. However, we keep optimistic that these problems can be solved if the potential of modern single-cell technologies integrated with other new methods are fully employed.

Based on the scRNA-seq data and knowledge of TF targets in the budding yeast *Saccharomyces cerevisiae*^18,39-43^, we also explored TF regulation at the single cell level and found distinctive temporal regulation of TFs during ageing in yeast. *YAP1* is a key TF responding to oxidative stress^44,45^. While it was highly activated in 16-hr/F compared to 16-hr/S early age subgroup, no significant difference of its activities were observed between 36-hr/F and 36-hr/S late age subgroups (Fig. 4b, c), implicating its vital role during early ageing, which in turn affects overall lifespan. In contrast, *RPN4*, the TF essential for proteasome biogenesis and RLS extension^46,47^, was only prominently activated in 36-hr/F compared to 36-hr/S late age subgroup, suggesting a dramatic loss of proteostasis in the late age and slow-dividing subgroup^49^ (Fig. 4b, c; Supplementary Fig. 8a-c; Supplementary Fig. 9a, b). These findings point not only to early but also late heterogeneity during ageing in yeast, and provide novel insights into understanding the molecular mechanisms of ageing that will lead to therapeutics for healthy ageing in humans ultimately^51^.

## Acknowledgement

We thank the BIOPIC sequencing platform at Peking University for the assistance of high-throughput sequencing experiments. This work was supported by Ministry of Science and Technology of China (2018YFA0108100), National Natural Science Foundation of China (21917802, 21525521), 2018 Beijing Brain Initiative (Z181100001518004), and Beijing Advanced Innovation Center for Genomics.

## Author contributions

Y.Z. and Y.H. conceived and designed the project. Y.Z., J.W., Y.S., S.J., X.Z., and G.K.A conducted the experiments. Y.Z., J.W., B.K., Q.L., J.W., X.Z. and Y.H. analyzed the data. Y.Z., J.W., B.K., X.Z., and Y.H. wrote the manuscript with the help from all other authors.

## Methods

### Strains and growth conditions

WT *Saccharomyces cerevisiae* in both BY4741 and BY4742 backgrounds were used for single-cell imaging analysis. The strain of *Hsp104*-GFP was derived from the standard GFP strain library in WT BY4741 background. WT BY4742 background was used in scRNA-seq during aging. WT BY4741 background was used in the replicative lifespan assay of *FIT3*Δ. For single-cell imaging, the cells were grown in the YPD liquid media before and after loading into the microfluidic chips. For scRNA-seq during aging and replicative lifespan assay of *FIT3*Δ, the cells were grown on SD solid agar plates.

### Single-cell imaging data analysis

The approach for single-cell imaging data analysis has been reported in detail elsewhere^11^. Yeast cell culture was grown in YPED at 30°C with OD600 of 0.5 before loading into the microfluidic device by a syringe connected to an automatically controlled peristaltic pump. The microfluidic device was mounted on a Nikon TE2000 time-lapsed microscope by a customized holder. Bright field images were taken once every 10 minutes throughout the whole life, and fluorescent images were taken once every 2 hours or 4 hours for measuring the *HSP104*-GFP level. The images were processed by ImageJ and MATLAB.

### Dissection and isolation of single cells for RNA-seq

We first inoculated WT yeast cells onto a solid agar plate with SD media and followed a standard protocol of replicative lifespan assay by continual (no storage in the 4°C fridge overnight) manual microdissection^15^. At 3 time points (2hr, 16hr and 36hr after birth), single yeast aging cells from the plate were manually dissected and placed individually into a single tube prefilled with lysis buffer containing zymolyase (3 × 10^−2^ U/µl) for efficiently digesting the cell wall and external RNA control consortium (ERCC) spike-in (8000 molecules) for assessing technical noise, followed by immediate freeze in liquid nitrogen and then storage in a -80°C freezer.

### Library preparation for scRNA-seq

After collecting all the single yeast aging cells, we performed scRNA-seq based on Smart-seq2^16,17^ with fine optimization. To efficiently lyse the single yeast aging cell and avoid possible mRNA degradation, we vigorously vortexed the lysis tubes for 1 min in a cold room. Then we kept the lysis tubes at 30°C for 10 min, followed by 3 min at 72°C. Subsequently, we added the RT reaction mix (RT-buffer and Invitrogen SuperScript II) for reverse transcription. Reverse transcription was carried out at 42°C for 90 min first, followed by 12 rounds of temperature cycling between 50°C and 42°C with 2 min each. The reaction was heat inactivated at 70°C for 15 min and then cooled down to 4°C. The oligo-dT and TSO primers used here were biotinylated to avoid potential production of excessive primer dimers and concatamers. After RT, the cDNA were amplified between 20∼25 cycles using KAPA HiFi enzyme. After cDNA amplification, the samples were purified using Agencourt AMPure XP beads at 0.8X bead concentration and quantified using Qubit Hs Assay (Life Technologies). We also checked the samples by a fragment analyzer to confirm the clean peak at ∼1.7 kb before subsequent processing. 1∼2 ng of cDNA was subjected to a tagmentation-based protocol (Vazyme TruePrep Kit) with 10 min at 55°C and dual index amplification for the library with 8∼12 cycles. The final libraries were purified twice using AMPure XP beads at 0.8X bead concentration and resuspended in 15∼20 µl elution buffer. Libraries were then quantified using Qubit Hs Assay before pooling for sequencing. Sequencing was performed in paired-end mode using Illumina NextSeq.

### scRNA-seq data pre-processing and filtering

Paired-end reads were mapped to the S288c *Saccharomyces cerevisiae* genome R64 version (www.yeastgenome.org) with ERCC spike-in sequences added using HISAT2 (version 2.1.0). Resulting bam files were sorted and indexed using samtools (version 1.1). Final read counts mapped to genes were extracted using FeatureCounts. Sequenced single yeast aging cells were removed from the analysis if they have < 1000 genes detected and 40,000 total mapped reads per cell, or if the proportion of ERCC spike-ins to total-mapped reads was > 0.74. After filtering, a scRNA-seq data set with 125 single yeast aging cells was used for the subsequent analysis.

### Normalization

Unless noted, normalization of raw read counts was done using the DESeq2^25^ package (v.1.22.2) in R. The size factor was computed by a formula embedded in DESeq2 for each cell based on the raw read counts matrix of all samples. Then these size factors were applied for normalizing different cells and finally the gene expression values are presented in the log_2_ space (log_2_NormCounts).

### Estimation of cell-to-cell transcriptional variability and identification of highly variable genes

We used two methods to estimate the cell-to-cell transcriptional variability during aging in yeast. The first was a correlation based method modified from Enge, M. *et al*^20^, where the transcriptional noise was expressed as biological variation over technical variation. First, we calculated the biological variation b_ij_ = 1-cor(x_ij_, u_i_), where u_i_ was the mean gene expression vector for the single cells in age group of i (2hr, 16hr and 36hr), and x_ij_ was the gene expression vector of cell j in the age group of i. Next, we calculated the corresponding technical variation t_ij_ = 1-cor(x^contr^_ij_, u^contr^) where x^contr^_ij_ and u^contr^ are the expression vector and mean expression vector of the ERCC spike-in controls. Finally the measurement of b_ij_/t_ij_ which reflected the biological noise as a fraction of technical noise for each cell was used for boxplot across different age groups as shown in Fig.1b. The second method was based on quantitative statistics reported previously^21^ (see Supplementary Note 6 of Brennecke et al^21^ for details of the statistical model). Briefly, to infer the genes that were highly variable within each age group, a linear regression model was applied to fit the relationship between the squared coefficient of variation (CV^2^) and the mean expression of ERCC spike-ins, and only genes with biological squared coefficient of variation > 0.25 (CV^2^ > 0.25) and FDR< 0.1 after multiple testing correction were regarded as HVGs.

### Differential gene expression and GO analysis

The differential gene expression analysis between pairwise age groups and subgroups was based on DESeq2^25^ with default parameters, taking log_2_FC >1 and adjusted P value < 0.05 as significant. GO analysis of these differentially expressed genes was performed by functional annotation tool of DAVID^22^ that classify the ontology of each gene into biological process or cellular component. The GO term enrichment results derived from DAVID were further verified alternatively by the R package of ClusterProfiler^27^.

### Weighted gene co-expression network analysis

WGCNA^28-29^ was performed on normalized gene expression data from DESeq2^25^, using 2498 genes, which are selected by removing unclassified genes (grey module) from the first round of WGCNA^28-29^. Then the second round WGCNA^29-30^ was performed following the standard process. Briefly, the topological overlap matrix (TOM) was constructed with a soft Power and was set to 2. The hub genes for each module were identified as module membership based group trait > 0.65 and gene significance > 0.2.

### Statistical analysis of regulation of transcription factor between age subgroups

To identify transcription factors with distinct regulation between age subgroups, 3 statistical approaches were applied stringently. The first one was to conventionally comparison of TF targets expression between age subgroups. The TF targets expression was defined as the averaged normalized expression of each set of TF targets for each cell. And we took log_2_FC (FoldChange) of median TF targets expression between age subgroups >1 (log_2_FC > 1) and a welch *t* test P value < 0.01 as significant, which resulted in 16 and 11 TF targets respectively that were significantly activated in the age subgroups of 16-hr/F and 36-hr/F compared to their counterparts (Supplementary Fig. 8a, b; Supplementary Table. 6). The second one was to further run a Wilcoxon rank sum test for each single cell that compare internally the normalized gene expression levels of each set of TF targets to all other detected genes for that cell, taking P < 0.0001 as criterion (indicated as regulon activity “on”), followed by intersection with TF targets derived from the first approach. This approach was similar with that from Gasch et al^18^. The last one was to correlate the TF targets expression with the generation of single cells in the age groups of 16-hr and 36-hr respectively, taking P < 0.05 as criterion, followed by intersection with TF targets derived from the former two approaches to avoid potential false positive results.

### PCA analysis

Raw read counts matrix with or without cell-cycle-regulated periodic genes^23^ were used as inputs for PCA by Seurat^52^. When the cell-cycle-regulated periodic genes were included, Seurat generates 631 common variable genes of all 125 single yeast aging cells, whose normalized read counts are further applied for PCA. When the cell-cycle-regulated periodic genes were excluded, Seurat generated 599 common variable genes of all 125 single yeast aging cells for PCA.

## Data availability

scRNA-seq data generated in this study has been uploaded to Gene Expression Omnibus under accession number xxxxxx.

## Figure legends

**Supplementary Fig**.**1** | **Early heterogeneity of cell divisions during ageing in yeast. a**, The distribution of generation at 8 hr, 12 hr and 16 hr after birth of single yeasts respectively (n=67). The plot showed the heterogeneity of cell divisions occurs early during ageing in yeast as indicated by the mean (Mean) and standard deviation (Std) of generation in the figure. **b**, The lifespan was plotted against the generation at 8 hr, 12 hr and 16 hr after birth of single yeasts respectively. It show a positive correlation. **c**, The *HSP104*-GFP level was plotted against the generation at 8 hr, 12 hr and 16 hr after birth of single yeasts respectively. It show a negative correlation. Each red dot in **b** or **c** represented a single cell with its generation and its final lifespan or *HSP104*-GFP level, while blue line was a linear fit with gray area indicating 0.95 confidence interval; correlation coefficient (R) and P value (P) were also shown.

**Supplementary Fig**.**2** | **Data filtering and technical assessment of scRNA-seq. a**, The number of raw read counts plotted against the number of genes detected per cell between different age groups. **b**, The ERCC ratio plotted against the number of genes detected per cell between different age groups. **c**, The ERCC ratio plotted against the number of raw read counts per cell between different age groups. Each dot in **a-c** represented a single cell with color indicating the age group or filtering status it belonged to (n=136 cells). **d** and **e** were mean normalized read counts and detection rate (the probability to have a read count number more than 0) plotted against the absolute number of RNA molecules per cell for each of the 92 ERCC RNA spike-in across all the single yeast aging cells that were filtered (n=125 cells).

**Supplementary Fig**.**3** | **Identification of HVGs within different age groups with or without cell-cycle-regulated periodic genes. a**, Squared coefficients of variation were plotted against the average normalized read counts for each cell within different age groups with cell-cycle-regulated periodic genes included. A gene was considered as HVG if the coefficient of biological variation was more than 0.5 (with the false discovery rate of 0.1). Red line represented the technical noise fit estimated by the ERCC spike-in RNA^21^ (see Methods). Endogenous genes, ERCC and HVGs were shown in black, green and magenta dots respectively. **b**, Venn diagrams of HVGs within different age groups and the putative cell-cycle-regulated periodic genes. The increased cell-to-cell transcriptional variability during ageing still existed even excluding these cell-cycle-regulated periodic HVGs from 3 age groups. **c**, PCA plot of single cells (n=125) from different age groups (with cell-cycle-regulated periodic genes included as input for PCA). The 3 age groups were segregated along the first PCA component successfully. **d**, Visualized plots of top 30 genes by absolute loading values for the first PCA component, with or without cell-cycle-regulated periodic genes included as input for PCA. **e**, Venn diagrams of the genes with top 30 genes by absolute loading values for the first PCA component, with or without cell-cycle-regulated periodic genes included as input for PCA, overlapped with putative cell-cycle-regulated periodic genes. **f**, Pseudotime trajectory of single cells (n=125) from different age groups (with cell-cycle-regulated periodic genes included as input). The youngest 2-hr age group was very concentrated, whereas the 16-hr early age group and 36-hr late age group were very scattered in order.

**Supplementary Fig**.**4** | **Differential gene expression between age groups. a**, Volcano plot of global differential gene expression analysis between different age groups using DESeq2 (see Methods). The criteria for statistical significance were log_2_ foldchange of absolute normalized gene expression more than 1 (Log_2_|FC|>1) and adjusted P value less than 0.05 (P_adj_<0.05). **b**, Boxplots of the average normalized expression of typical upregulated and downregulated gene categories identified in the early age group of 16hr compared to 2hr, across different age groups. Each black dot in **b** represented a single cell. *p < 0.05 and ****p < 1.4 × 10^−10^ from Wilcoxon rank sum test.

**Supplementary Fig**.**5** | **GO enrichment analysis between age groups. a-f** were GO enrichment analysis of differentially expressed genes from the pairwise comparison of 3 age groups using the R package clusterProfiler^27^ (see Methods). The number of genes in the enriched GO category was indicated by the size of the dot while the adjusted P value was indicated by the color of the dot.

**Supplementary Fig**.**6** | **GO analysis of hub genes of ageing related co-expression gene module identified by WGCNA. a**, GO terms of 52 hub genes of turquoise module. These hub genes were upregulated during ageing in yeast and are mainly enriched in OSR, oxidation-reduction process and even longevity regulating pathway. **b**, GO terms of 70 hub genes of blue module. These hub genes were downregulated during ageing in yeast and are mainly enriched in ribosome biogenesis.

**Supplementary Fig**.**7** | **Correlation of gene expression and the generation of single cells in the early and late age groups. a-c**, Normalized gene expression of *HAC1, FET3* and *FIT2* plotted against the generation of single cells in the 16-hr early age group and 36-hr late age group, respectively. Each red dot represented a single cell with the respective normalized gene expression and its generation, while blue line was a linear fit with gray area indicating 0.95 confidence interval; correlation coefficient (R) and P value (P) were also shown. They all showed negative correlation with statistical significance (P<0.05) except the *FIT2* at 16hr (P=0.14). **d**, Pearson correlation of normalized gene expression with the generation of single cells in the early age group of 16hr, taking P<0.05 as significant. The biological process of iron transport was enriched as negatively correlated while the ribosome biogenesis positively correlated. **e**, Boxplots of the average normalized expression of gene category of ribosome biogenesis identified in **d**, across different age groups and subgroups. Each black dot in **e** represented a single cell. ***p < 2.7 × 10^−4^, ****p < 3.3 ×10^−6^, and “ns” means not significant, from Wilcoxon rank sum test. **f**, Pearson correlation of gene expression with the generation of single cells in the 36-hr late age group, taking P<0.05 as significant. The biological process of iron transport was enriched as negatively correlated while the translation, mitochondrial translation and glycolytic process positively correlated.

**Supplementary Fig**.**8** | **Distinct regulation of TF between age subgroups. a**, 16 TF targets that were significantly activated in the early age subgroup of 16-hr/F by conventional comparison of median TF targets expressions to 16-hr/S, taking Log_2_FC>1 and P <0.01 as significant. **b**, 11 TF targets that were significantly activated in the late age subgroup of 36-hr/F by conventional comparison of median TF targets expressions to 36-hr/S, taking Log_2_FC>1 and P <0.01 as significant. **c**, The significantly activated TF targets in 16-hr/F and 36-hr/F in contrast to their counterparts were further narrowed down to 5 and 2 (highlighted in red in **a** and **b**) respectively by Wilcoxon rank sum test comparing normalized gene expression levels of each set of TF targets to that of all other detected genes for each cell, taking P < 0.0001 as the criterion and indicated as “on” of the regulon activity. **d** and **e** are boxplots of differential expression of 4 other TF targets that were highly expressed in the early age subgroup of 16-hr/F compared to 16-hr/S identified by the first two stringent statistical approaches (see Methods), across different age groups and subgroups, respectively. Each black dot in **d** and **e** represented a single cell. *p < 0.05, **p < 0.01, ***p < 2.8 × 10^−4^, ****p < 7.1 × 10^−5^, and “ns” means not significant, from Wilcoxon rank sum test.

**Supplementary Fig**.**9** | **Correlation of TF targets expression with the generation of single cells in the early and late age groups. a**, Pearson correlation of median TF targets expression with the generation of single cells in the 16-hr early age group, taking P<0.05 as significant. The expression of *YAP1* targets was found to be most positively correlated. **b**, Pearson correlation of median TF targets expression with the generation of single cells in the 36-hr late age group, take P<0.05 as significant. The expression of 2 *RPN4* targets identified by previous two statistical approaches also positively correlated with the generation of single cells in the 36-hr late age group.

