## Supplementary Materials for "Single-cell RNA-seq reveals early heterogeneity during ageing in yeast"

### Figure S1, Zhang et al.

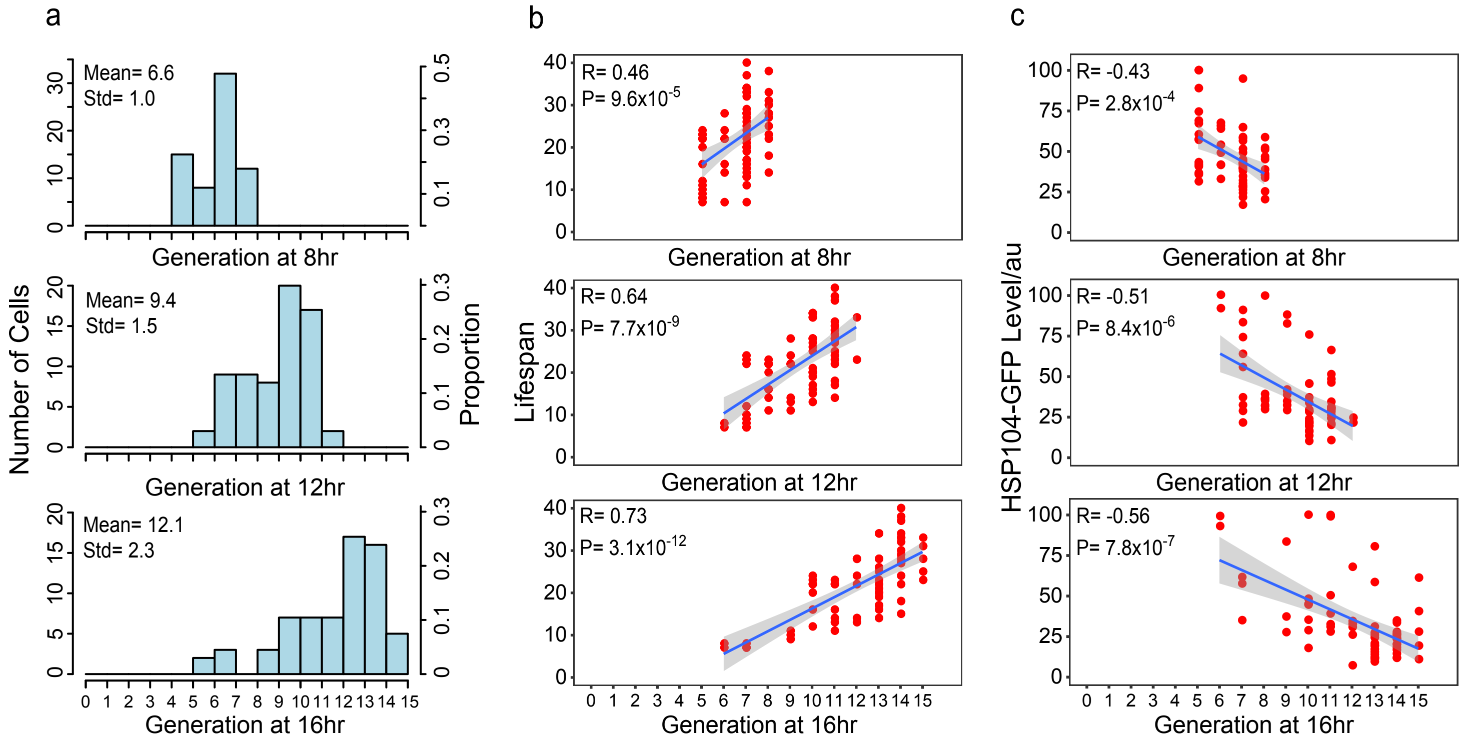

Figure S2, Zhang et al.

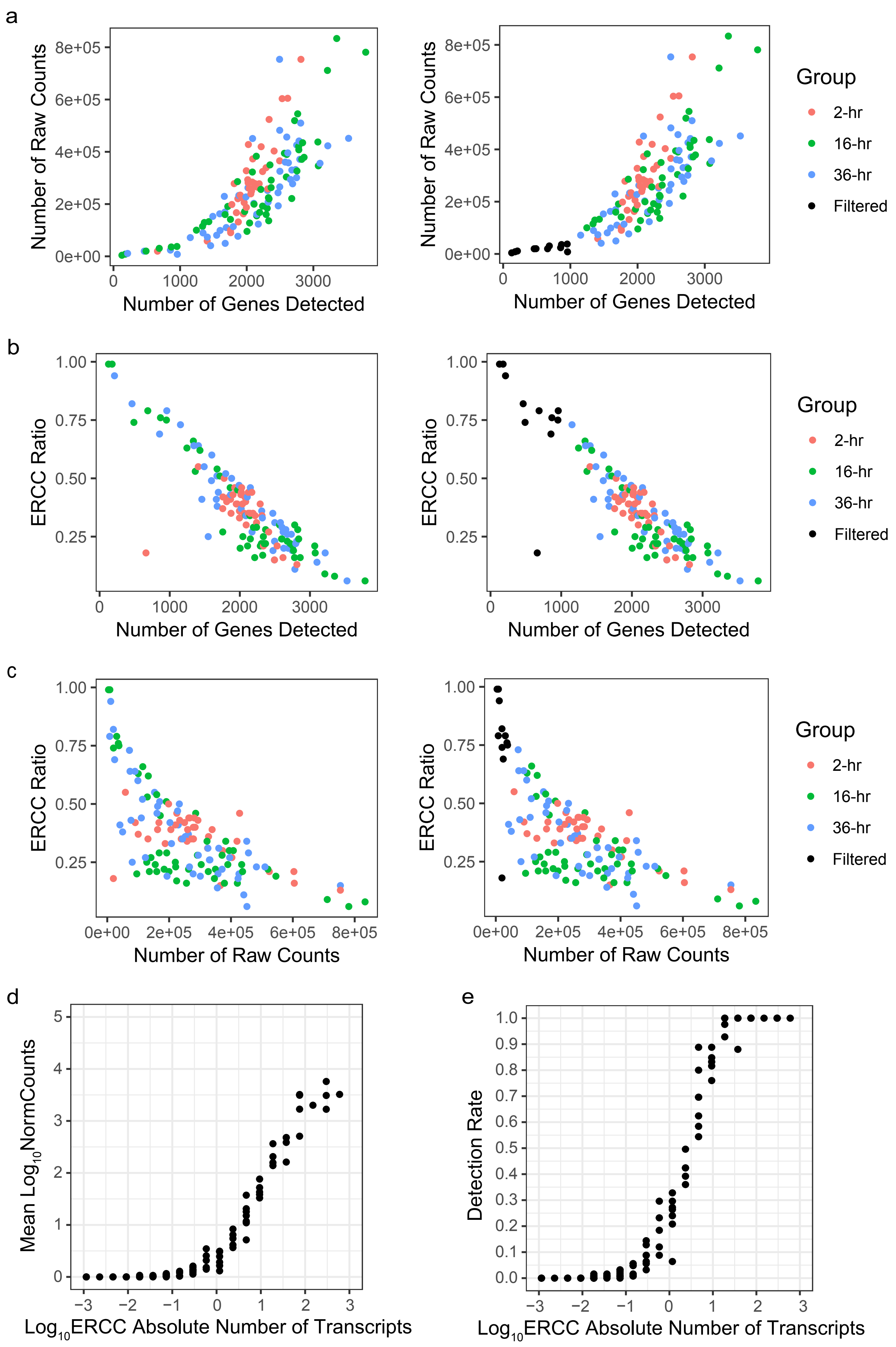

Figure S3, Zhang et al.

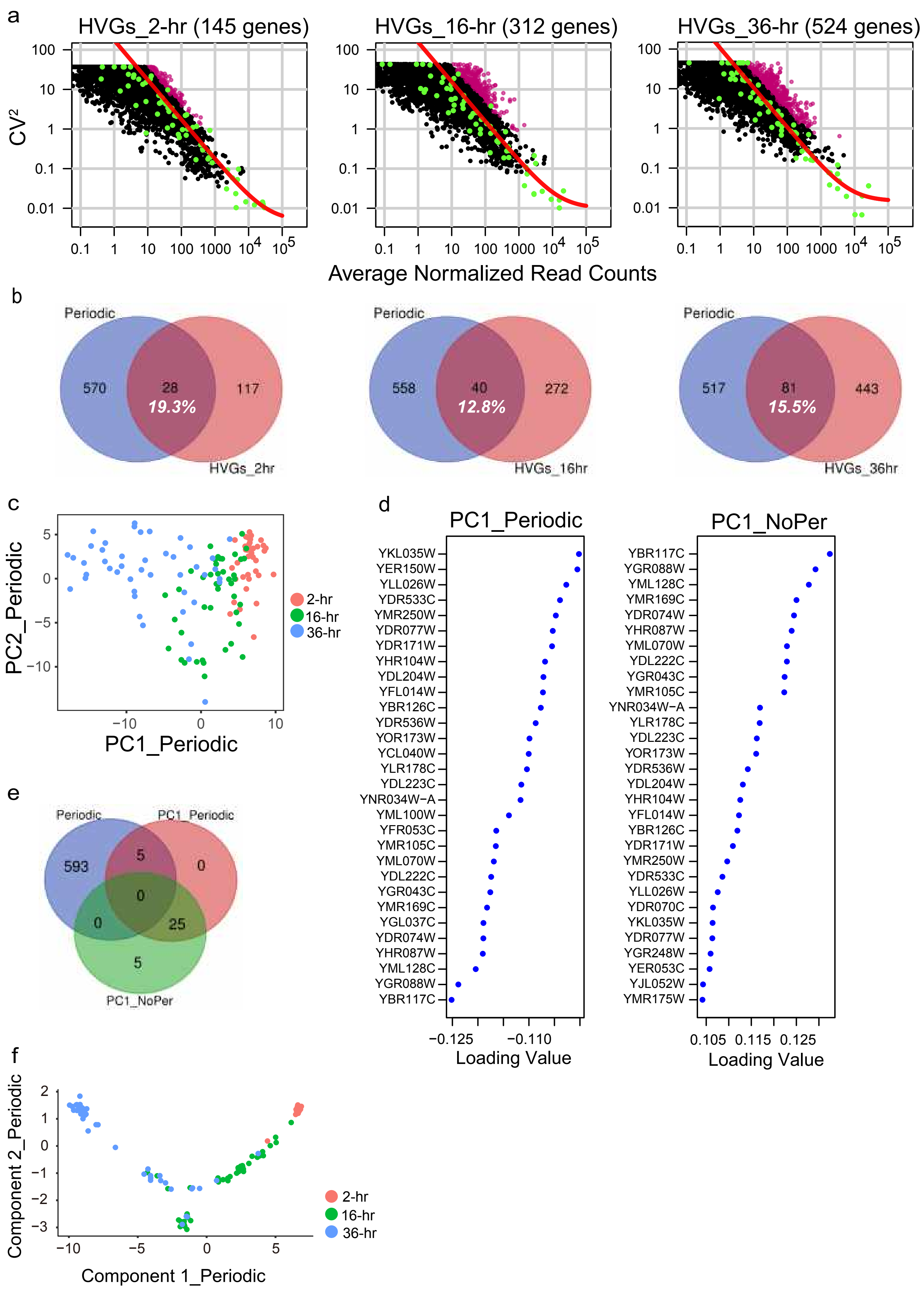

Figure S4, Zhang et al.

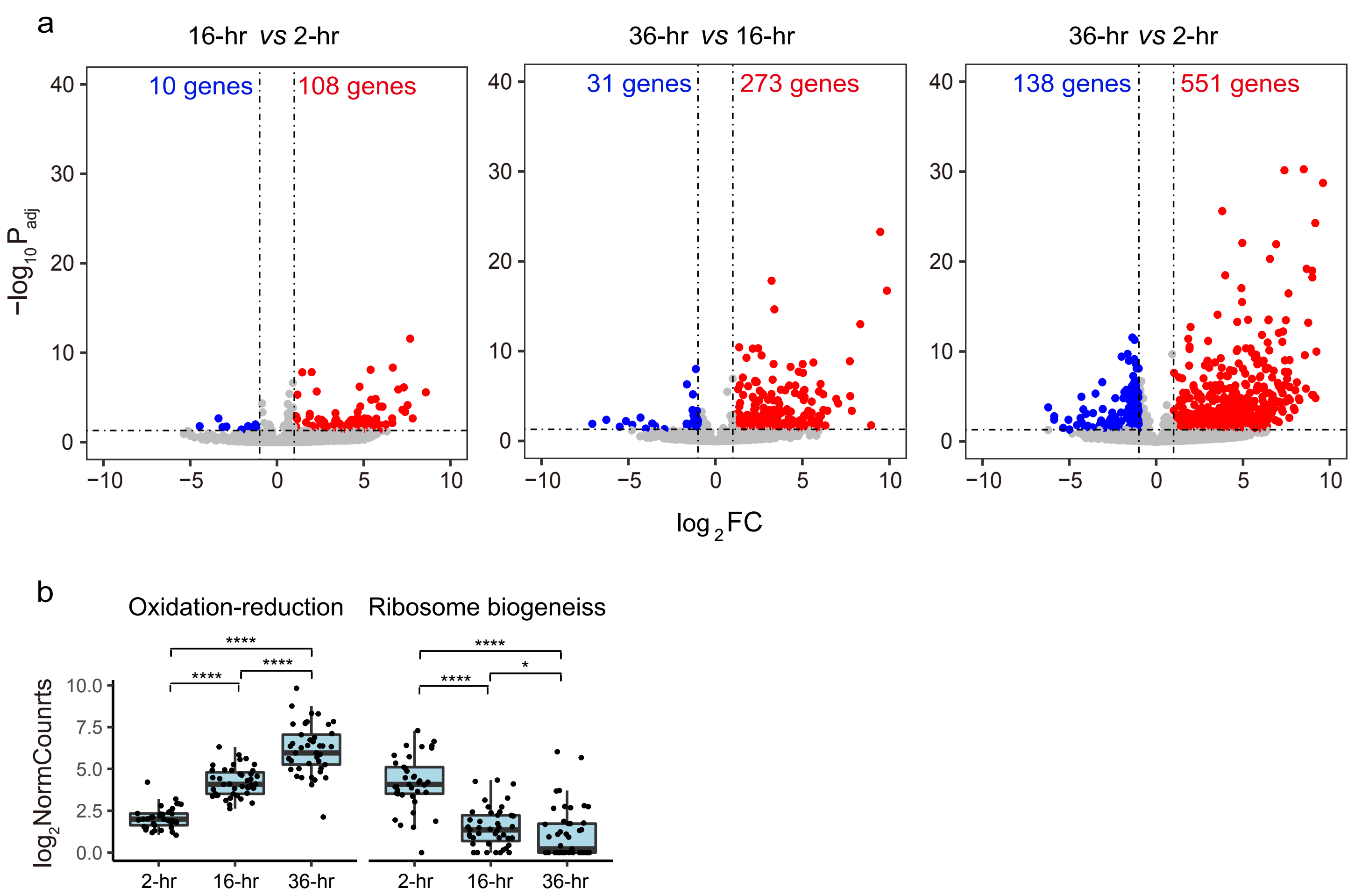

### Figure S5, Zhang et al.

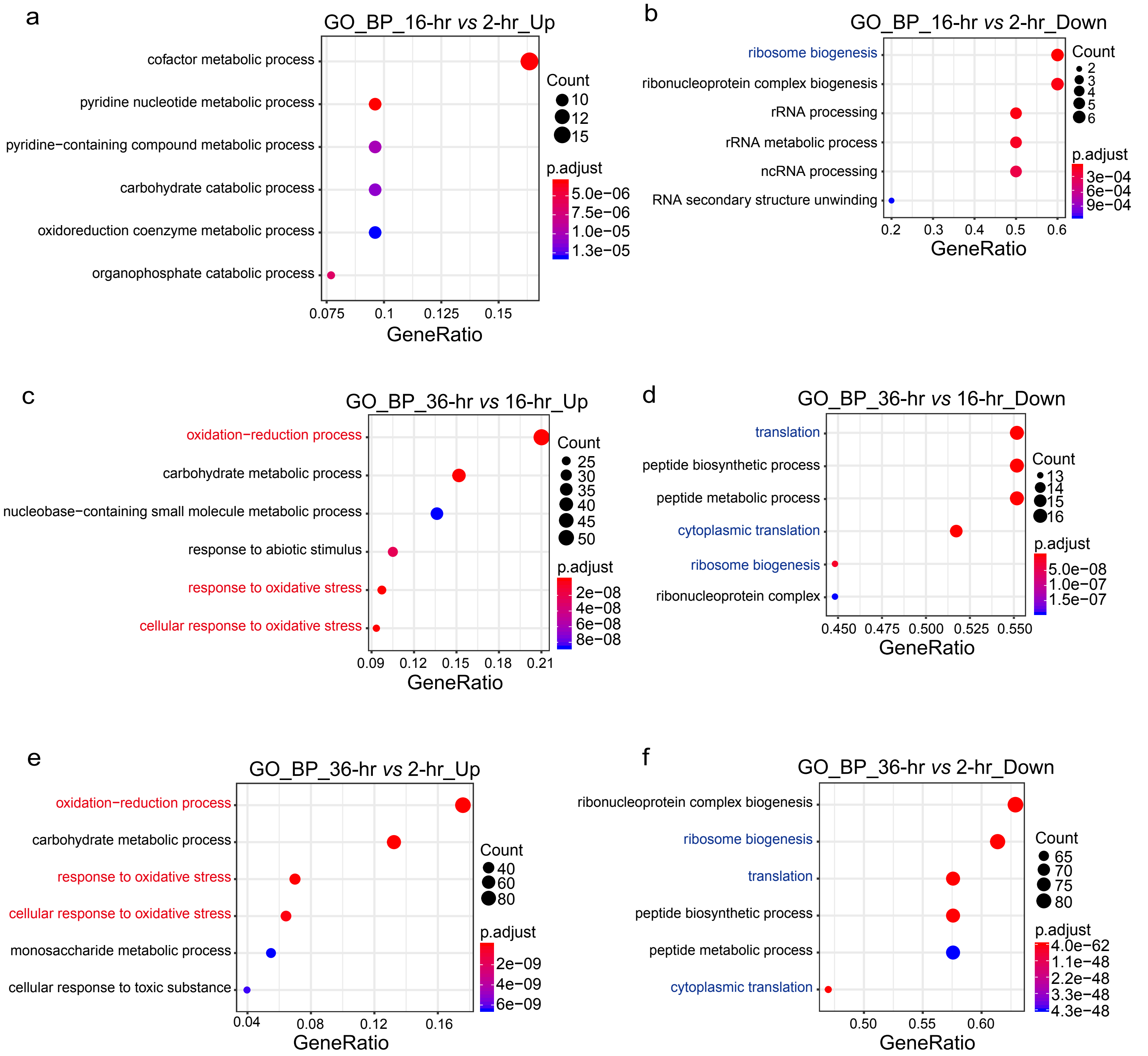

Figure S6, Zhang et al.

a

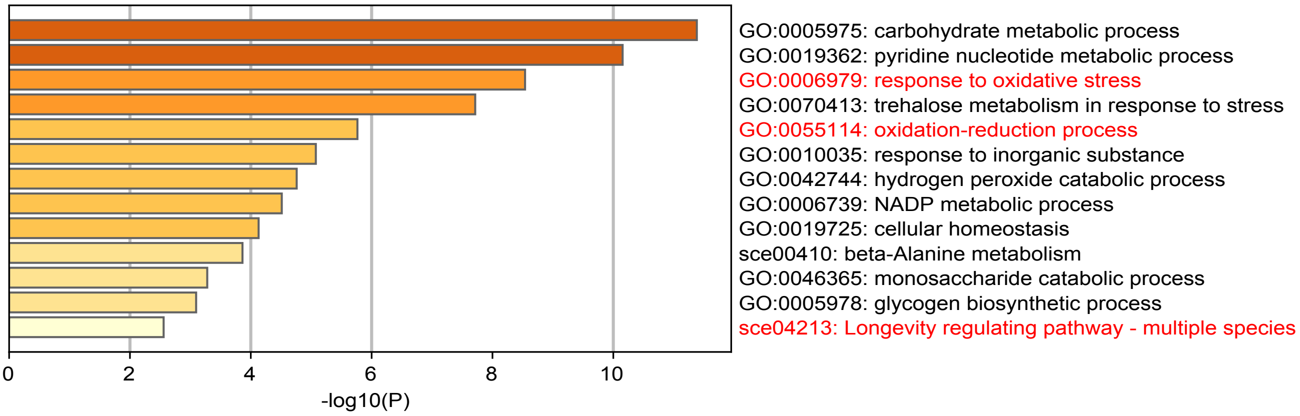


b

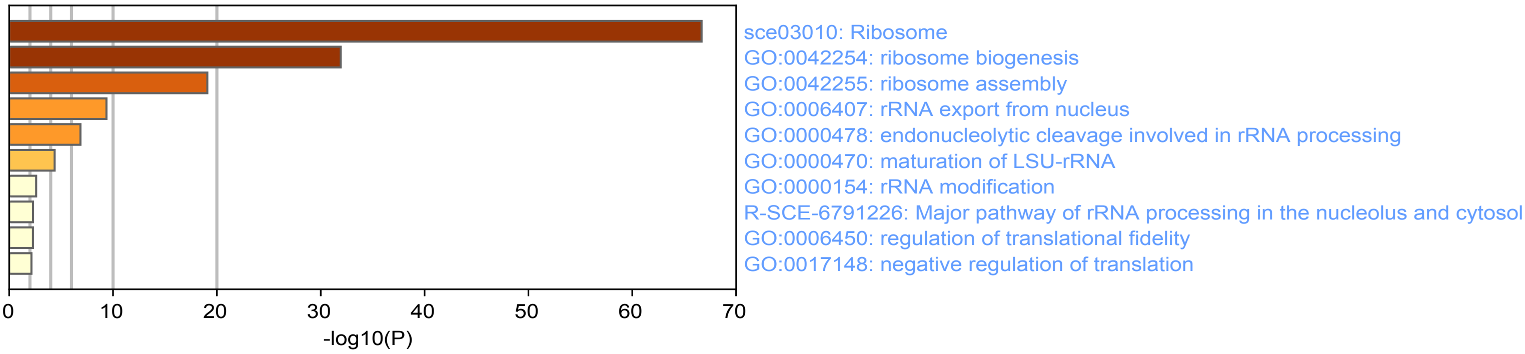

Figure S7, Zhang et al.

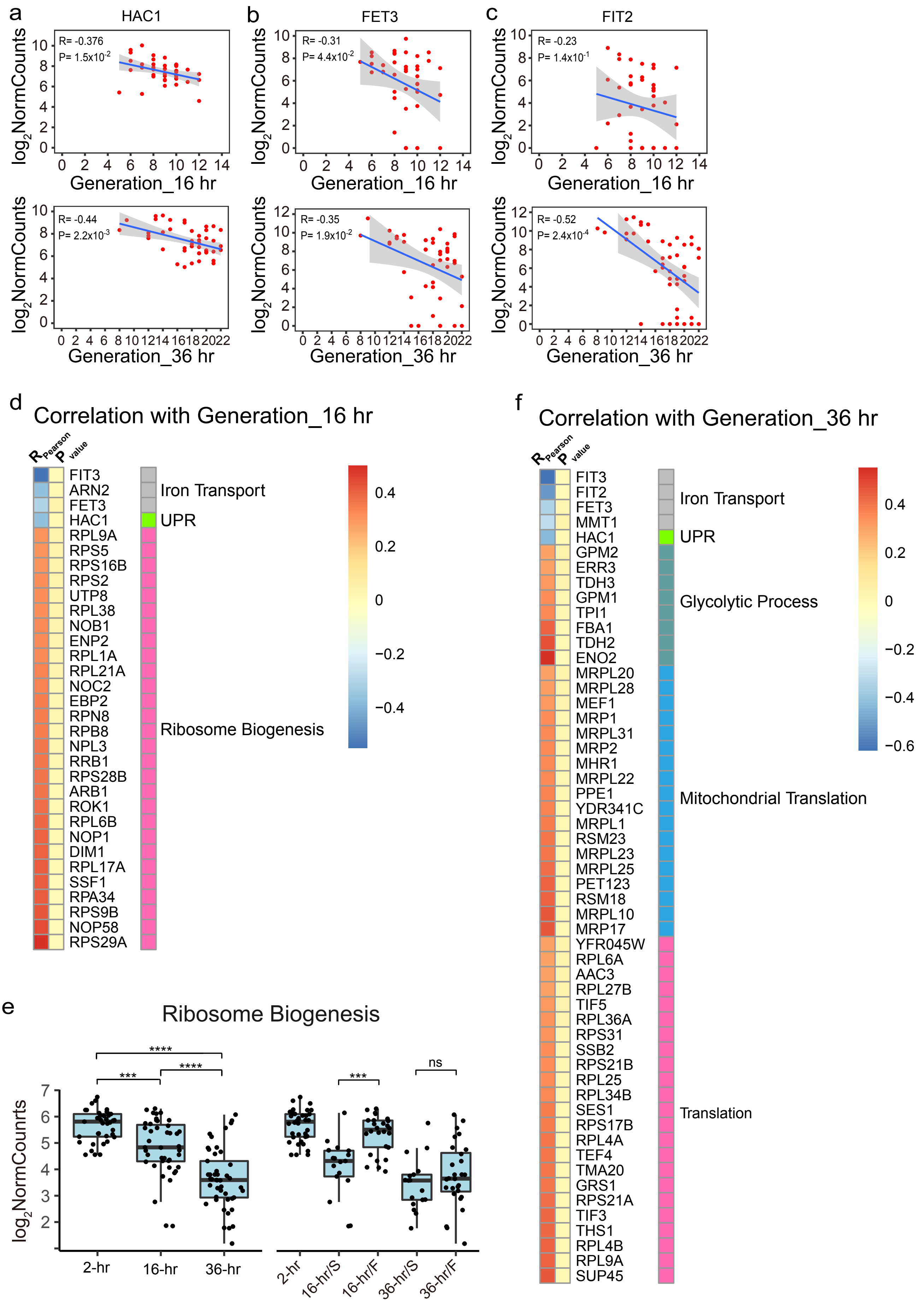

Figure S8, Zhang et al.

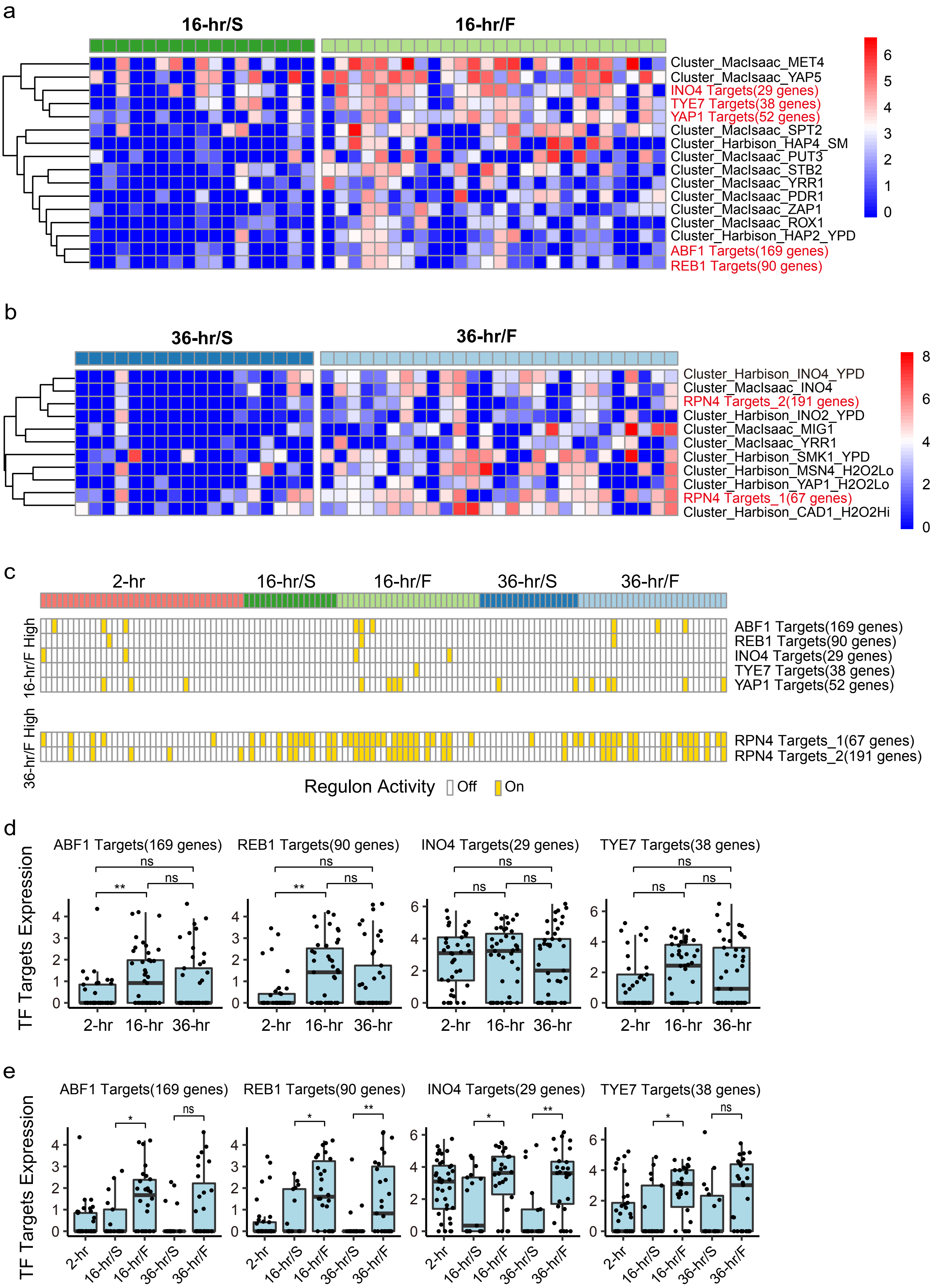

Figure S9, Zhang et al.

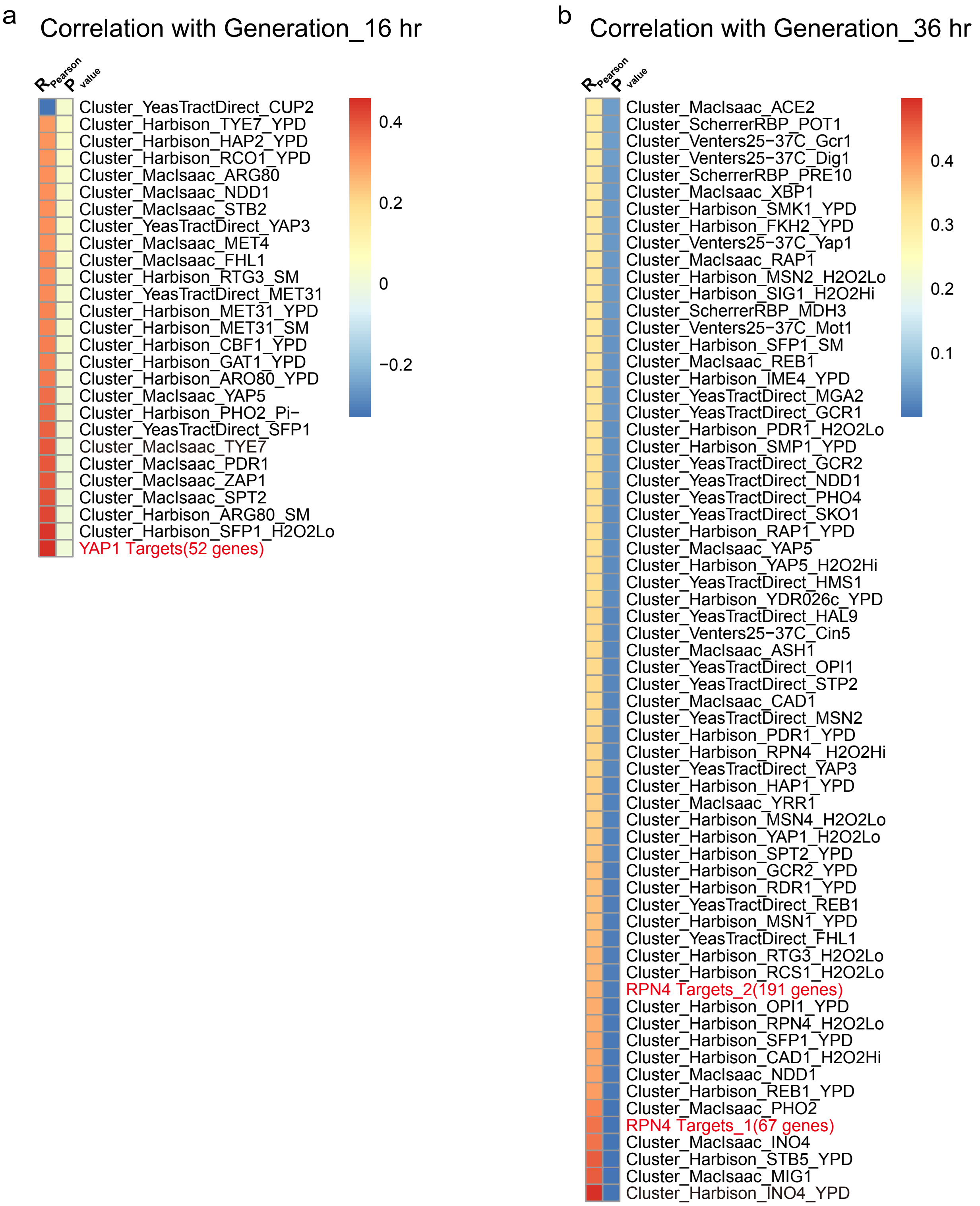
